# Hybrid genome assembly and evidence-based annotation of the egg parasitoid and biological control agent *Trichogramma brassicae*

**DOI:** 10.1101/2020.04.06.027094

**Authors:** K. B. Ferguson, T. Kursch-Metz, E. C. Verhulst, B. A. Pannebakker

## Abstract

*Trichogramma brassicae* (Bezdenko) are egg parasitoids that are used throughout the world as biological control agents and in laboratories as model species. Despite this ubiquity, few genetic resources exist beyond COI, ITS2, and RAPD markers. Aided by a *Wolbachia* infection, a wild-caught strain from Germany was reared for low heterozygosity and sequenced in a hybrid *de novo* strategy, after which several assembling strategies were evaluated. The best assembly, derived from a DBG2OLC-based pipeline, yielded a genome of 235 Mbp made up of 1,572 contigs with an N50 of 556,663 bp. Following a rigorous *ab initio*-, homology-, and evidence-based annotation, 16,905 genes were annotated and functionally described. As an example of the utility of the genome, a simple ortholog cluster analysis was performed with sister species *T. pretiosum*, revealing over 6000 shared clusters and under 400 clusters unique to each species. The genome and transcriptome presented here provides an essential resource for comparative genomics of the commercially relevant genus *Trichogramma*, but also for research into molecular evolution, ecology, and breeding of *T. brassicae*.

## INTRODUCTION

The chalcidoid *Trichogramma brassicae* (Bezdenko) (Hymenoptera: Trichogrammatidae) is a minute parasitoid wasp (∼0.5 mm in length) that develops within the eggs of other insects (Smith, 1996). For over 50 years, it has been in use world-wide as a biological control agent as many lepidopteran pests of different crops are suitable hosts (Polaszek, 2009). The most common application of *T. brassicae* in Europe is against *Ostrinia nubilalis* (Hubner) (Lepidoptera: Pyralidae), the European corn borer. For example, in 2003 alone, over 11000 ha of maize in Germany was treated with *T. brassicae* (Zimmermann, 2004). It is also released against lepidopteran pests in spinach fields as well as in greenhouses (e.g. tomato, pepper, and cucumber) (Klug and Meyhöfer, 2009). With its wide application in biological control, *T. brassicae* is a well-studied species. Field trials have been conducted on several aspects, such as host location and dispersal behaviour (Suverkropp et al., 2010, 2009), overwintering ability (Babendreier et al., 2003), while other biological control related studies considered issues related to low temperature storage (Lessard and Boivin, 2013), reaction to insecticides (Delpuech and Delahaye, 2013; Ghorbani et al., 2016; Jamshidnia et al., 2018; Liu and Zhang, 2012; Thubru et al., 2018), or risk assessment (Kuske et al., 2004).

Next to its application as a biological control agent, this tiny parasitoid has been used in other research, both in genetic studies (Cruaud et al., 2018; Laurent et al., 1998; Wajnberg, 1993) and ecological studies (Cusumano et al., 2015; Fatouros and Huigens, 2012; Huigens et al., 2009). In addition, several initiatives investigate the infection of *T. brassicae* with *Wolbachia* bacteria (Ivezić et al., 2018; Poorjavad et al., 2012) and the consequences of such an infection (Farrokhi et al., 2010; Poorjavad et al., 2018; Rahimi-Kaldeh et al., 2018). As *T. brassicae* is a cryptic species with several other congenerics, misidentification and misclassification is a known issue (Polaszek, 2009). In response, molecular identification of trichogrammatids is well studied and established (Ivezić et al., 2018; Rugman-Jones and Stouthamer, 2017; Stouthamer et al., 1999; Sumer et al., 2009). Recently, several RADseq libraries were constructed from single *T. brassicae* wasps to aide in resolving the aforementioned phylogenetic issues within *Trichogramma* (Cruaud et al., 2018). Otherwise, the genomics of *T. brassicae* have largely been neglected even though a well annotated genome would allow researchers and biological control practitioners access to a wealth of information and open new avenues for comparative genomics and transcriptomics for evolutionary, ecological, and applied research.

Here, we report the whole-genome sequencing and annotation of a *T. brassicae* strain infected by *Wolbachia* that had thelytokous reproduction, in which females arise from unfertilized eggs. A hybrid *de novo* sequencing strategy was chosen to address two common issues: we used long PacBio Sequel reads to bridge the large segments of repetitive sequences often found in Hymenoptera, while countering the error bias of long read technology with the accuracy of Illumina short reads. A similar strategy was recently applied to improve the *Apis mellifera* genome, where the long PacBio reads were the backbone that boosted the overall contiguity of the genome, alongside the incorporation of repetitive regions (Wallberg et al., 2019).

In this report, we present the hybrid *de novo* genome of *T. brassicae*. Three different assemblers were evaluated, and the most complete genome assembly was used for decontamination and *ab initio*-, homology-, and evidence-based annotation. The resulting annotation was functionally described using gene ontology analysis. Finally, a heterozygosity comparison and simple ortholog cluster analysis with the congeneric *T. pretiosum* was performed, which can be considered a starting point for future comparative genomics of the commercially important genus *Trichogramma*.

## METHODS

### Species origin and description

Individuals of *Trichogramma brassicae* were acquired by AMW Nützlinge GmbH (Pfungstadt, Germany). The strain was baited in May 2013 in an apple orchard near Eberstadt, Germany. The orchard was surrounded by blackberry hedges, forest, and other orchards. For baiting, the eggs of *Sitotroga cerealella* (Olivier) (Lepidoptera: Gelechiidae) (Mega Corn Ltd., Bulgaria) were glued on paper cards (AMW Nützlinge GmbH, Germany), usually used for releasing *Trichogramma* sp. in corn fields and households. These cards were placed directly into the trees, approximately two meters above ground. After five days in the field, baiting cards were collected and incubated together at 25°C. Following emergence, individuals were kept together, offered *S. cerealella* eggs, and reared in a climate chamber (27±2°C, L:D=24:0h for four days, then transferred to16±2°C, L:D=0:24h until emergence).

In 2016, the offspring of twenty isolated females were transferred to Wageningen University (The Netherlands) to be reared for low heterozygosity. The resulting offspring were reared in a single general population on irradiated *Ephestia kuehniella* (Zeller) (Lepidoptera: Pyralidae) eggs as factitious hosts under laboratory conditions in a climate chamber (20 ± 5°C, RH 50 ± 5%, L:D=12:12 h). *Wolbachia* presence was determined following the PCR amplification protocol of Zhou et al 1998 in a presence/absence assessment with known positive and negative control samples (Zhou et al., 1998). Natural *Wolbachia* infections have previously been detected in Iranian populations of *T. brassicae* (Farrokhi et al., 2010), but none of the Eurasian populations have been known to support this symbiosis (Stouthamer, 1997; Stouthamer and Huigens, 2003).

### Isofemale line

Following confirmation of *Wolbachia* infection (Supplementary materials S1.1.1), a single female from the general population was isolated (generation 0, G0), and given eggs *ad libitum*. In the resulting generation (G1), unmated females were isolated and reared with eggs *ad libitum*. Offspring of the initial isolations G0 and G1 were confirmed to be entirely female, suggesting thelytokous parthenogenetic reproduction. Combined with isolating single females, this maximizes genetic similarity of the following generation (G2) of these G1 females. One of these G2 strains, S301, was boosted for multiple generations over the period of one year. By the time of collection for sequencing, both the S301 and general population no longer harboured *Wolbachia* at detectable levels (Supplementary materials S1.1.2).

### gDNA extraction

Three separate extractions were prepared in 1.5 mL safelock tubes with each several hundred *Trichogramma brassicae*. The tubes were frozen in liquid nitrogen with approximately six 1-mm glass beads and shaken for 30 s in a Silamat S6 shaker (Ivoclar Vivadent, Schaan, Liechtenstein). DNA was then extracted using the Qiagen MagAttract Kit (Qiagen, Hilden, Germany). Following an overnight lysis step with Buffer ATL and proteinase K at 56°C, extraction was performed according to the MagAttract Kit protocol. Elutions were performed in two steps with Buffer AE (Tris-EDTA) each time (first 60 µL, then 40 µL), yielding 100 µL. The two extractions yielding the largest amount of DNA (5.49 µg and 8.24 µg) were combined for long-read sequencing, while the remaining extraction (1.67 µg) was used for short-read sequencing. DNA concentration was measured with an Invitrogen Qubit 2.0 fluorometer using the dsDNA HS Assay Kit (Thermo Fisher Scientific, Waltham, USA) while fragment length was confirmed on gel.

### Library preparation and sequencing

Sequence coverage was calculated using the previously established genome size estimate for *T. brassicae* of 246 Mbp (Johnston et al., 2004). Library preparation and sequencing was performed by Novogene Bioinformatics Technology Co., Ltd., (Beijing, China). For Illumina sequencing, gDNA was used to construct one paired-end (PE) library according to the standard protocol for Illumina with an average insert size of 150 bp and was sequenced using an Illumina HiSeq 2000 (Illumina, San Diego, USA). For Single Molecule Real Time (SMRT) sequencing, gDNA was selected for optimal size using a Blue Pippin size selection system (Sage Science, Beverley, USA) following a standard library preparation. The library was then sequenced on a PacBio Sequel (Pacific Biosciences, Menlo Park, USA) with 16 SMRT cells.

### Assembly and decontamination

Prior to assembly, Illumina reads were assessed for quality using *FASTQC* (Andrews et al., 2015), then trimmed for quality in *CLC Genomics Workbench 11* using default settings (Qiagen). Trimmed Illumina reads were paired for subsequent analysis.

In order to achieve the best possible assembly, three assembly pipelines were evaluated: one for PacBio-only reads and two hybrid assemblers. The PacBio-only were assembled with *Canu* (v1.6) with modifications based on PacBio Sequel reads (correctedErrorRate=0.085 corMhapSensitivity=normal alongside corMhapSensivity=normal) (Koren et al., 2017). This is assembly version v1.0 in the subsequent discussion.

The first hybrid assembly pipeline using both long and short sequencing read sets was *SPAdes* (v3.11.1) (Bankevich et al., 2012). The *SPAdes* genome toolkit supports hybrid assemblies with the *hybridSPAdes* algorithm (Antipov et al., 2016). Three iterations of the *SPAdes* pipeline were run with varying k-mer sizes resulting in three different assembly versions: 21, 33, 55 (default, v2.1); k-mer sizes 21, 33, 55, 77 (v2.2); and a single k-mer size of 127 (v2.3).

The second hybrid assembly pipeline was *DBG2OLC* (Ye et al., 2016). The *DBG2OLC* pipeline can be readily tweaked with other programs depending on the job (Chakraborty et al., 2016). Following the *DBG2OLC* pipeline, de Bruijn graph contigs were generated using *SparseAssembler* using default settings and setting the expected genome size to 750 Mbp to ensure a genome size output that is unrestricted (Ye et al., 2012). Contigs were transformed into read overlaps using *DBG2OLC* with settings suggested for large genomes and PacBio Sequel data (k=17; AdaptiveTh=0.01; KmerCovTh=2; MinOverlap=20; RemoveChimera=1), according to the *DBG2OLC* manual (https://github.com/yechengxi/DBG2OLC). This creates an assembly backbone of the best overlaps between the short-read de Bruijn contigs and the long reads. *minimap2* (v2.9) and *Racon* (v1.0.2) were used for consensus calling remaining overlaps to the assembly backbone (Li, 2018; Vaser et al., 2017). The resulting consensus assembly was polished twice using the Illumina reads with *Pilon* (v1.22) (Walker et al., 2014). This final assembly is v3.0 in subsequent discussion.

The best of the five assemblies generated was determined on the basis of N50, genome size, and completeness (Table 1). Genome statistics such as N50, number of contig, and genome size were determined using *Quast* (Gurevich et al., 2013). Assembly completeness was assessed using *BUSCO* (v3.0.2) with the insect_odb9 ortholog set and the fly training parameter (Simão et al., 2015). Based on these characteristics, the decision was made to move forward with assembly v3.0, which was then decontaminated for microbial sequences using NCBI *BLASTn* (v2.2.31+) against the NCBI nucleotide collection (nr).

**Table 1.**
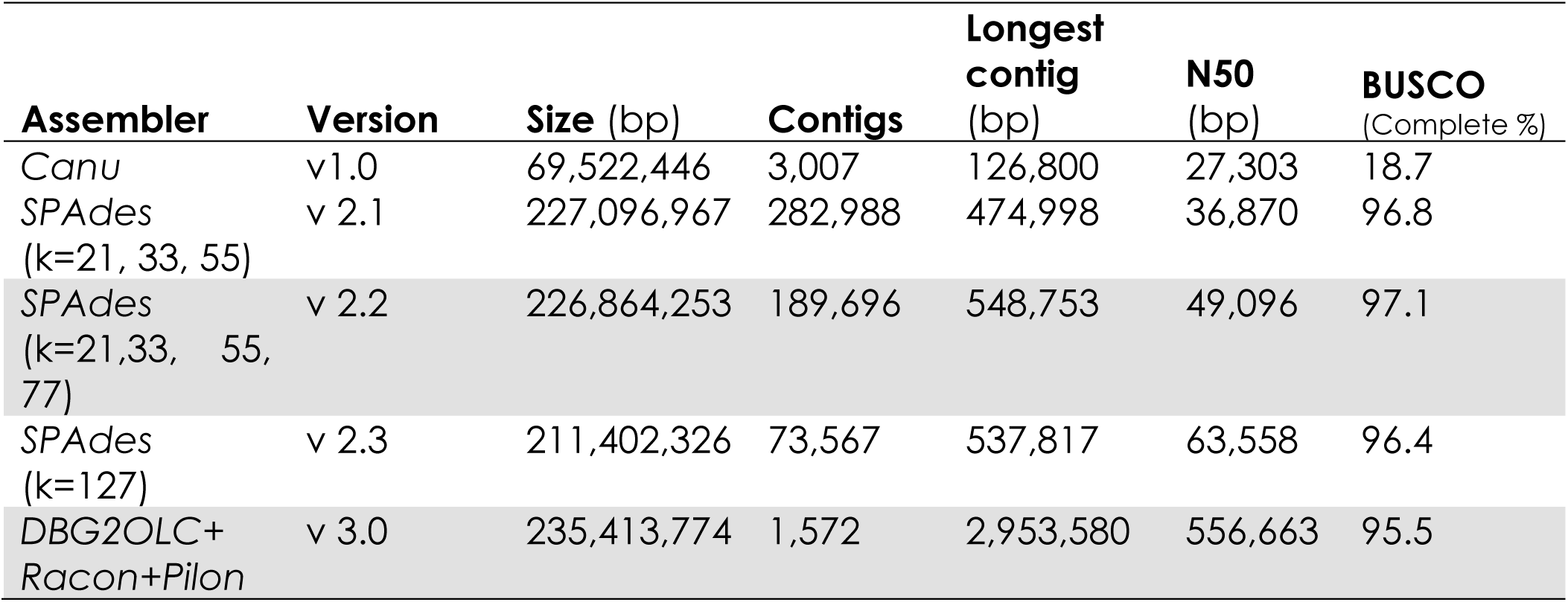
Statistics for five assemblies of *Trichogramma brassicae*. The first strategy was PacBio-only in *Canu*, while three hybrid assembly strategies were based on *SPAdes* and modulating k-mer sizes, and an additional hybrid assembly was based on an adapted *DBG2OLC*+*Racon*+*Pilon* protocol. BUSCO score is based on the <u>insect_db09 dataset (Simão et al., 2015).</u>

### *Wolbachia* contamination

Two contigs contained a large amount of *Wolbachia* content, with over 80% of the scaffold containing material with 75% or higher homology to *Wolbachia*. These contigs were assessed for homology against the NCBI nucleotide collection (nr) and removed from the assembly (Supplementary material S1.2). Post-decontamination, the assembly is referred to as v3.5.

### RNA extraction, library construction, and sequencing

*T. brassicae* wasps from the S301 line were collected for RNAseq for evidence-based annotation. Hundreds of adult individuals (male and female) were killed by freezing at −80°C, then frozen in liquid nitrogen in a single 1.5 mL safelock tube with approximately six 1-mm glass beads and shaken for 30 s in a Silamat S6 shaker (Ivoclar Vivadent). The RNeasy Blood and Tissue Kit (Qiagen) was used according to manufacturer’s instructions, and final column elution was achieved using 60 µL sterilized water. The sample was measured for quality and RNA quantity using an Invitrogen Qubit 2.0 fluorometer and the RNA BR Assay Kit (Thermo Fisher Scientific). The RNA sample was then processed by Novogene Bioinformatics Technology Co., Ltd., (Beijing, China) using poly(A) selection followed by cDNA synthesis with random hexamers and library construction with an insert size of 300 bp. Paired-end sequencing was performed on an Illumina HiSeq 4000 according to manufacturer’s instruction. Quality filtering was applied to remove adapters, reads with more than 10% undetermined bases, and reads of low quality for more than 50% of the total bases (Qscore less than or equal to 5).

### *Ab initio* gene finding, transcriptome assembly, and annotation

For the *ab initio* gene finding, a training set was established using the reference genome of *Drosophila melanogaster* (Meigen) (Diptera: Drosophilidae) (Genbank: GCA_000001215.4; Release 6 plus ISO1 MT) and the associated annotation (Adams et al., 2000; Dos Santos et al., 2015). The training parameters were used by *GlimmerHMM* (v3.0.1) for gene finding in the *T. brassicae* genome assembly v3.5 (Majoros et al., 2004). For homology-based gene prediction, *GeMoMa* v1.6 was used with the *D. melanogaster* reference genome alongside our RNAseq data as evidence for splice site prediction (Keilwagen et al., 2016). For evidence-based gene finding, the pooled RNAseq data was mapped to the to the *T. brassicae* genome separately with *TopHat* (v2.0.14) with default settings (Trapnell et al., 2009). After mapping, *Cufflinks* (v2.2.1) was used to assemble transcripts (Trapnell et al., 2010). *CodingQuarry* (v1.2) was used for gene finding in the genome using the assembled transcripts, with the strandness setting set to ‘unstranded’ (Testa et al., 2015).

The tool *EVidenceModeler (EVM)* (v1.1.1) was used to combine the *ab initio*, homology-based, and evidence-based information, with evidence-based weighted 1, *ab initio* weighted 2, and homology-based weighted 3 (Haas et al. 2008). We annotated the predicted proteins with *BLASTp* (v2.2.31+) on a custom database containing all SwissProt and Refseq genes of *D. melanogaster* (Acland et al., 2014; Boutet et al., 2008; Camacho et al., 2009), followed by an additional search in the NCBI non-redundant protein database (nr) to obtain additional homology data.

### GO term analysis

A list of genes was constructed for Gene Ontology (GO) term classification by deduplicating the annotated proteins and removing the non-annotated proteins. These accession IDs were converted into UniProtKB accession IDs using the UniProt ID mapping feature and deduplicated a final time (Boutet et al., 2008). These UniProtKB accession IDs were in turn used with the *DAVID 6*.*8 Functional Annotation Tool* to assign GO terms to each accession ID with the *D. melanogaster* background and generate initial functional analyses (Huang et al., 2009a, 2009b) (see supplementary S1.3 for *DAVID* input list).

### Heterozygosity estimates

The heterozygosity of the S301 line was assessed using sequence reads and k-mer counting, and compared to the congeneric *Trichogramma pretiosum* (Riley) (Hymenoptera: Trichogrammatidae), for which sequence data exists for both a thelytokous (asexual) *Wolbachia*-infected strain as well as an inbred arrhenotokous (sexual) line (Lindsey et al., 2018). Using *jellyfish* (v2.3.0) to count k-mers, the same trimmed and paired Illumina reads used for assembly were assessed using the default k-mer size of 21 (m=21), with results exported to a histogram (Marçais and Kingsford, 2011). This histogram file was then used with *GenomeScope* (v1.0) to estimate heterozygosity of the reads based on a statistical model, where a Poisson distribution is expected for a homozygous sample while a bimodal distribution is expected for a homozygous distribution (Vurture et al., 2017). This genome profiling gives a reliable estimate for heterozygosity as well as estimates of repetitive content. The same *jellyfish* and *GenomeScope* analyses were performed on *T. pretiosum* short-read sequence data for the thelytokous strain (NCBI SRA database, SRR1191749) and the arrhenotokous line (SRR6447489), with adaptions for reported insert sizes (Lindsey et al., 2018).

### Ortholog cluster analysis

The complete gene set of *T. brassicae* was compared to that of *T. pretiosum* (Lindsey et al., 2018), which was retrieved from the i5K Workspace (Poelchau et al., 2016). An ortholog cluster analysis was performed on both gene sets via *OrthoVenn2* with the default settings of E-values of 1e-5 and an inflation value of 1.5 (Xu et al., 2019). For *T. brassicae* protein set, see supplementary materials S1.5.

### Data availability

All sequence data are available at the EMBL-ENA database under BioProject PRJEB35413, including assembly (CADCXV010000000.1). An additional, complete annotation file (.gff) is also available (Ferguson, 2020). Additional data, such as gel images, the *Wolbachia* contaminated contigs, input gene list for *DAVID, GenomeScope* images, and complete protein set are available via the supplementary materials.

## RESULTS AND DISCUSSION

### Sequencing, assembly, and decontamination

Sequencing of the Illumina 150 bp paired-end library yielded 80,489,816 reads. After quality filtering and trimming, 80,483,128 paired-end reads were retained. Sequencing the PacBio Sequel library yielded 2,500,204 subreads with an average length of 6377 bp. The genome size estimate for *T. brassicae* is 246 Mbp (Johnston et al., 2004) indicating that short-read coverage was 98x while long-read coverage was 64x, resulting in a total coverage of 162x. Three assembly pipelines were used, resulting in five potential assemblies where one, v3.0, was eventually selected for further use. Results of these assemblies are detailed in Table 1.

The first draft assembly generated with Canu with the altered settings for PacBio Sequel data resulted in an assembly of approximately 70 Mbp in size, drastically smaller than the 246 Mbp expected, and contained a total of 3,007 contigs with an N50 of 27,303. The longest contig was 126,800 bp in size.

The second assembly strategy relied on hybrid assembly pipelines, and *SPAdes* was used with the default k-mer settings, which resulted in an assembly of approximately 227 Mbp in size with an N50 of 36,870 and a *BUSCO* completeness of 96.8%. Three different assembly runs were done with differing k-mer sizes: the default k-mer sizes of 21, 33, 55 (v2.1); default k-mer sizes plus 77 (v2.2); or the highest possible k-mer size of 127 (v2.3). Increasing the k-mer size only improved N50 scores to a point, along with decreasing the number of contigs, and stable *BUSCO* scores, however, the assembled genome size drops dramatically with the third attempt shrinking down to 211 Mbp. Based on *BUSCO* scores and N50 alone, the second *SPAdes* attempt, v2.2, would be the best of the three, though all three are similar in most measures.

The third assembly strategy used the *DGB2OLC*+*Racon*+*Pilon* pipeline, which resulted in assembly v3.0. Here, there is a large difference compared to the previous *SPAdes* assemblies. Particularly, the number of contigs is reduced dramatically from the 70,000 to 280,000 range of the *SPAdes* output down to a mere 1,572. Meanwhile, the assembled genome size is now 235 Mbp and with an N50 of 556,663 and a BUSCO score of 95.5%. The full completeness score for this assembly, using the 1658 BUSCO groups within the insect_od09 BUSCO set, returned 1531 (92.3%) complete and single-copy BUSCOs, 53 (3.2%) complete and duplicated BUSCOs, 22 (1.3%) fragmented BUSCOs, and 52 (3.2%) missing BUSCOs (Simão et al., 2015).

While the PacBio-only assembly in *Canu* could have been improved using different settings or additional tools, we decided to focus on using the additional sequence information of the Illumina reads in the subsequent hybrid assembly strategies. The *SPAdes* assemblies (v2.1-3) were already decent but could have been further improved using *Pilon*, a tool that improves assemblies at the base pair level using high quality Illumina data. However, the v3.0 assembly was by far the best assembly based on assembled genome size, N50, and BUSCO scores and therefore we chose this strategy for our *T. brassicae* genome assembly.

Decontamination of this assembly (v3.0) resulted in the removal of two contigs as the homology analysis using *BLASTn* with the NCBI nr database indicated that both contigs were confirmed to be largely composed of *Wolbachia* genomic content. Contig “Backbone_1176” is 9,448 bp in length and two areas of the contig, representing over 80% of its length, showed high homology to *Wolbachia*. Similarly, contig “Backbone_1392” is 17,350 bp and three separate areas representing over 80% showed similar levels of homology to *Wolbachia* After decontamination this final assembly (v3.5) was used for annotation.

### *Ab initio* gene finding, transcriptome assembly, and annotation

In our RNA sequencing experiment, we generated 26,479,830 150bp paired-end cDNA reads. Filtering the reads for quality retained 99.3% of these reads to be used for evidence-based gene finding via transcriptome assembly.

The annotations from the evidence-based gene finding were used alongside homology-based findings and *ab initio* annotations in a weighted model, resulting in a complete annotation for the assembly. In 865 mRNA tracks, representing approximately 5.1% of the official gene set, a gene model could not be annotated via the SwissProt database, and these tracks are named “No_blast_hit.” The majority of tracks are annotated with reference to SwissProt or GenBank accession number of the top *BLASTp* hit.

Transcriptome assembly and mapping resulted in 45,876,158 mapped transcripts (48,327,134 total). *CodingQuarry* predicted 45,454 evidence-based genes from these mapped transcripts, while *ab initio* gene finding using *GlimmerHMM* resulted in 16,877 genes and homology-based gene finding with *GeMoMa* resulted in 6,675 genes. The final complete gene set was created using *EVidenceModeler*, where a weighted model using all three inputs resulted in a complete gene set of 16,905 genes.

### GO term analysis

The complete gene set of 16,905 genes was deduplicated and genes with no correlating *BLASTp* hit were removed from this analysis. The remaining 9,373 genes were subjected to UniProtKB ID mapping, resulting in 8,247 genes with a matching ID after another round of deduplication (828 duplicates found). The remaining 755 accession IDs were not able to be matched, half of which are obsolete proteins within the UniParc database (377).

The *DAVID Functional Annotation Tool* used 6,585 genes for the analysis and showed that 80.8% (5,320) contribute to 530 biological processes, 77.5% (5,104) contribute to 115 different cellular component categories, and 74.2% (4,889) contribute to 93 molecular functions (genes can code to multiple GO terms). The remaining 1,662 genes are uncategorised.

### Heterozygosity estimates

Using short-read data and k-mer counting, heterozygosity was estimated for our isofemale S301 line and compared to both a parthenogenesis inducing *Wolbachia*-infected strain and an arrhenotokous line of *T. pretiosum* (Lindsey et al., 2018). The average estimated heterozygosity for our S301 *T. brassicae* line is 0.0332% with approximately 0.608% repetitive content (for full details, see Table 2). This is similar to the thelytokous *T. pretiosum* line, which has a slightly lower estimated heterozygosity (0.0289%) and a lower amount of repetitive content (0.482%). Both have a very distinct Poisson distribution, indicating a low heterozygosity (Figure S1.4.1-2). The arrhenotokous *T. pretiosum* showed a higher estimated heterozygosity (0.863%), a larger amount of repetitive content (2.64%), and a slightly bimodal distribution (Figure S1.4.3). The fact that both thelytokous *Trichogramma* species have a similar low level of heterozygosity when compared to the arrhenotokous *T. pretiosum* suggests that in both cases *Wolbachia* infection had a severe effect on genetic diversity. As the canonical mechanism of parthenogenesis-induction in other *Wolbachia* infected thelytokous *Trichogramma* species is gamete duplication (Pannebakker et al., 2004; Stouthamer and Kazmer, 1994), in which unfertilized eggs are diploidized and results in fully homozygous progeny in a single generation, the low genomic heterozygosity rate suggests a similar mechanism for *Wolbachia*-induced parthenogenesis in *T. brassicae*. However, the involvement of *Wolbachia* in causing all-female offspring in this *T. brassicae* strain and the presence and mechanisms of *Wolbachia* in other thelytokous *T. brassicae* strains (Farrokhi et al., 2010; Poorjavad et al., 2018, 2012) does require further investigation.

**Table 2.**
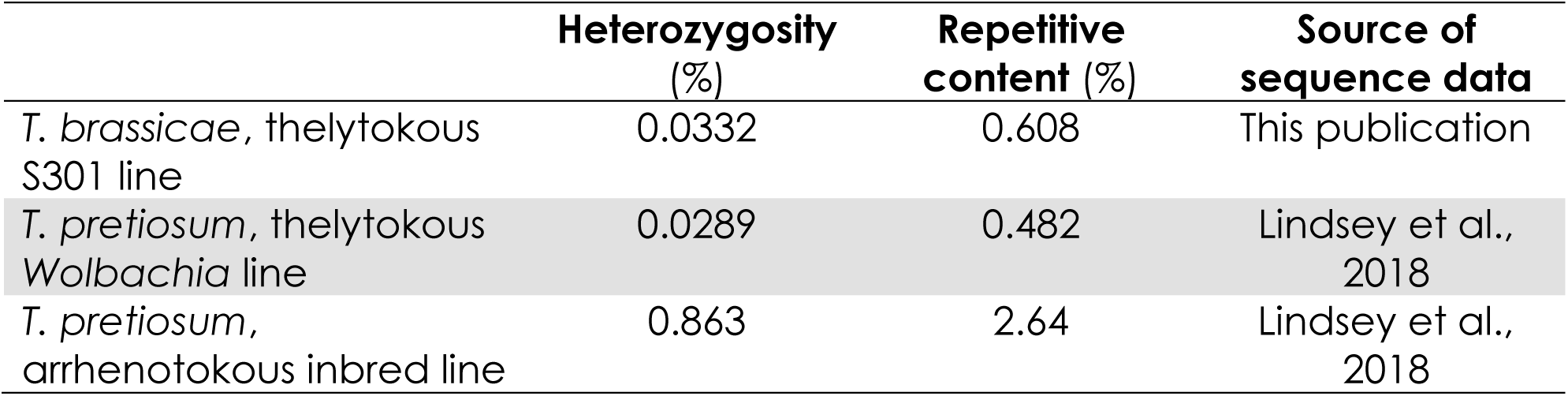
Heterozygosity and repetitive content analysis of *Trichogramma brassicae* (thelytokous), *Trichogramma pretiosum* (thelytokous), and *T. pretiosum* (arrhenotokous) lines based on sequence data.

|  | **Heterozygosity**<br>**(%)** | **Repetitive**<br>**content (%)** | **Source of**<br>**sequence data** |
| --- | --- | --- | --- |
| *T. brassicae*, thelytokous<br>S301 line | 0.0332 | 0.608 | This publication |
| *T. pretiosum*, thelytokous<br>*Wolbachia* line | 0.0289 | 0.482 | Lindsey et al.,<br>2018 |
| *T. pretiosum*,<br>arrhenotokous inbred line | 0.863 | 2.64 | Lindsey et al.,<br>2018 |

### Ortholog cluster analysis

The complete gene set of *T. brassicae* was compared to that of *T. pretiosum* using *OrthoVenn2* (full output in Table 3). Both species have a similar range of proteins (16,905 in *T. brassicae* and 13,200 in *T. pretiosum*) that form a similar number of clusters (6,537 in *T. brassicae* and 6,489 in *T. pretiosum*). The two species share 6,158 clusters (of 16,899 proteins), while *T. brassicae* has 379 unique clusters (1,726 proteins) and *T. pretiosum* has 331 unique clusters (1,005 proteins), as shown in Figure 1. These unique clusters account for approximately 5% of the entire cluster set for both species, and may both indicate true areas of differentiation, or result from differences in the annotation strategies. There is a similar amount of singleton clusters (proteins that do not cluster with others) in *T. brassicae* (5,291) and *T. pretiosum* (5,184). Both the unique clusters and the unique single-copy genes could be novel proteins, regions of contamination, evidence of unique horizontal gene transfer, or pseudogenes. More investigation into these protein clusters in addition to a more comprehensive manual annotation should shed some light on the differences between these closely related yet geographically distinct parasitoid wasps.

**Table 3.** Output of OrthoVenn2 ortholog cluster analysis of *Trichogramma brassicae* and *Trichogramma pretiosum*.

| Species | Proteins | Clusters | Singletons | Source of gene set |
| --- | --- | --- | --- | --- |
| *T. brassicae* | 16,905 | 6,537 | 5,291 | This work (S1.5) |
| *T. pretiosum* | 13,200 | 6,489 | 5,184 | Lindsey et al., 2018;<br>Poelchau et al., 2015 |

**Figure 1.**
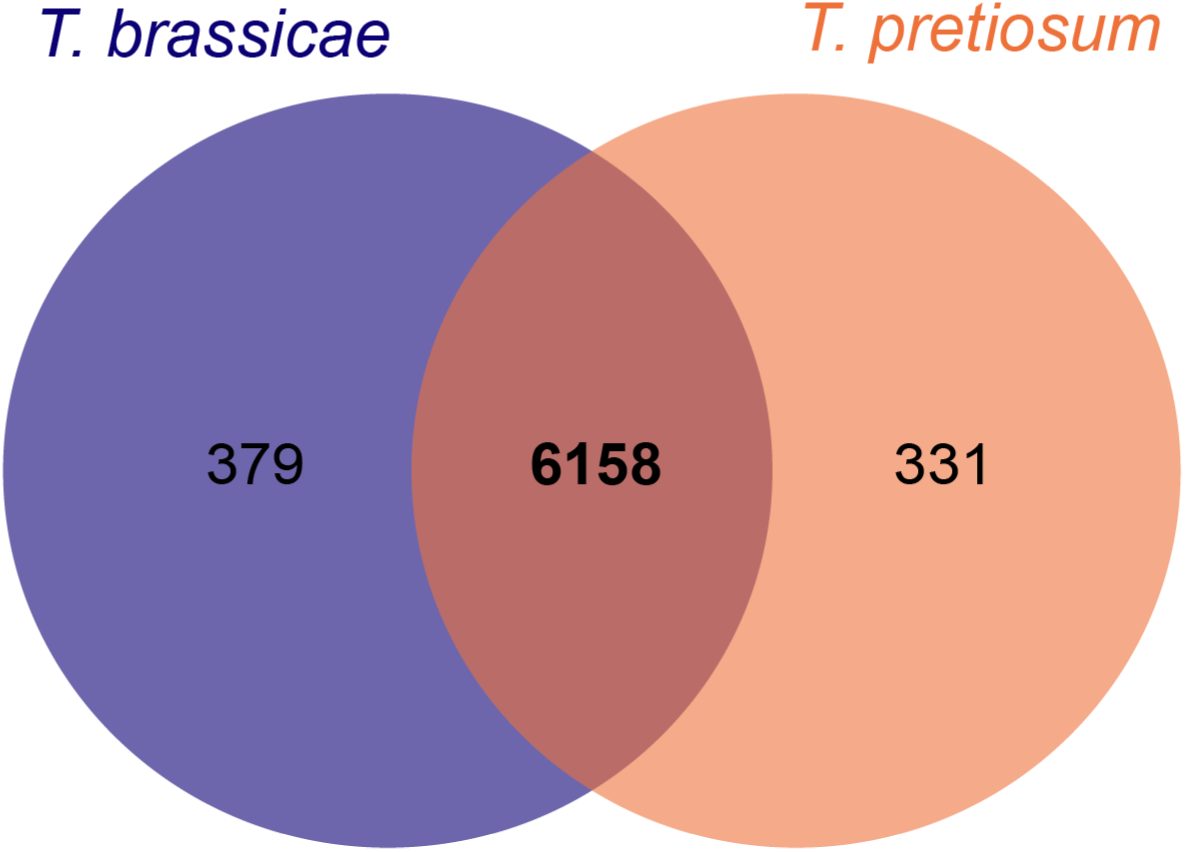
Ortholog clusters analysis between *Trichogramma brassicae* and *T. pretiosum* using OrthoVenn2 (Xu et al., 2019). The number of clusters shared between the two organisms is in bold.

## CONCLUSIONS AND PERSPECTIVES

Here, we present the genome of biological control agent *Trichogramma brassicae*, a chalcidoid wasp used throughout the world for augmentative biological control as well as genetic and ecological research. This unique strain hosted a parthenogenesis-inducing *Wolbachia* infection and is the first European *Trichogramma* genome to be published, allowing for comparative analyses with other *Trichogramma* genomes, as we have shown. Our genomic data also illuminates the possible mechanism of parthenogenesis-induction by *Wolbachia* in this strain. Furthermore, the variety of genomic and transcriptomic data generated for this genome provide much-need resources to bring *T. brassicae* into the -omics era of biological research.

A hybrid approach was used, resulting in a highly contiguous assembly of 1,572 contigs and 16,905 genes based on *ab initio*, homology-based, and evidence-based annotation, for a total assembly size of 235 Mbp. Two scaffolds were identified that were of *Wolbachia* origin and removed. Ortholog cluster analysis showed 379 unique protein clusters containing 1,726 proteins. Future studies are needed to show whether these clusters are truly unique. This genome and annotation provides the basis for future, more in-depth comparative studies into the genetics, evolution, ecology, and biological control use of *Trichogramma* species.

## ACKNOWLEDGEMENTS AND FUNDING

We would like to thank Bernd Wührer (AMW Nützlinge) for providing access to specimens; Gabrielle Bukovinszkine Kiss, José van de Belt, Frank Becker (Wageningen University), and Lorraine Latchoumane for assistance with rearing, DNA, and RNA extraction; Richard Stouthamer and Nina Fatouros for sharing their *Trichogramma* knowledge; and Sophie Chattington, Andra Thiel (University of Bremen), and Bas Zwaan (Wageningen University) for their assistance in this project. GenomeScan B. V. performed decontamination and annotation analysis. This work has received funding from the European Union Horizon 2020 research and innovation programme under the Marie Sklodowska-Curie grant agreement no. 641456.

## SUPPLEMENTARY MATERIALS

Additional supplementary material from this study is available in an attached document, with some material found on the DANS EASY Repository, https://doi.org/10.17026/dans-23w-a9tn (explanation within supporting document).

