## supplementary materials for "Hybrid genome assembly and evidence-based annotation of the egg parasitoid and biological control agent *Trichogramma brassicae*"

Material accessible on DANS EASY:

Pannebakker, dr. ir. B. A. (Wageningen University); Ferguson, K. B. (Wageningen University) (2019): The *Trichogramma brassicae* genome, supporting data. DANS. https://doi.org/10.17026/dans-23w-a9tn

### S1.1 *Wolbachia* detection

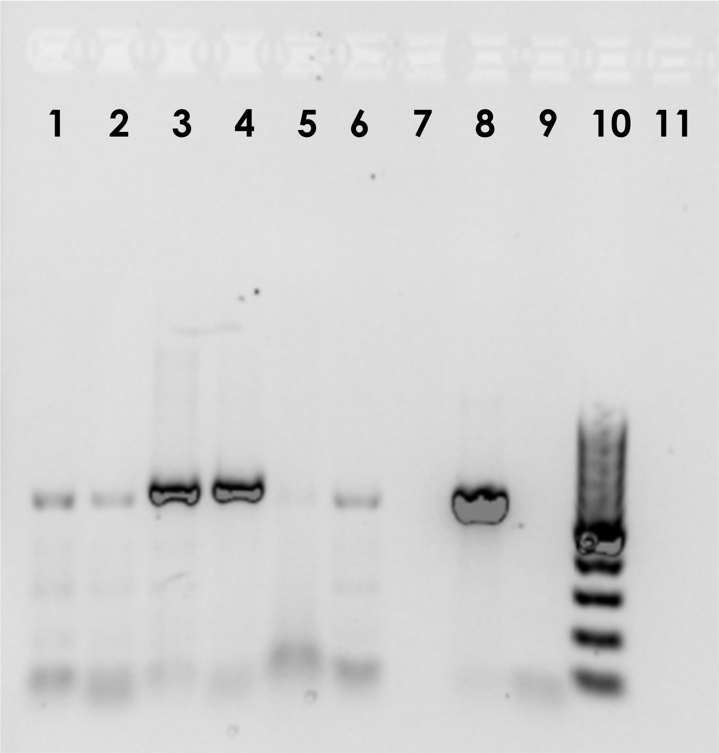

#### Figure S1.1.1 Initial *Wolbachia* detection via PCR

*Wolbachia* detection was carried out on isolated DNA from individual males and female wasps, as well as population samples of several individuals, according to the PCR protocol of (Zhou et al., 1998). Lanes 1 and 2 are male *T. brassicae*, 3 and 4 are female *T. brassicae*, and 5 and 6 are pooled populations from the general population of *T. brassicae*. Lane 7 was a negative PCR control (sterilised water). Lane 8 is positive *Wolbachia* control *Muscidifurax uniraptor* (Hymenoptera: Pteromalidae) from a laboratory population confirmed to have thelytoky-inducing *Wolbachia* (Gottlieb and Zchori-Fein, 2001). Lane 9 is negative *Wolbachia* control *Nasonia vitripennis* from the *Wolbachia* cured AsymCx line (refer to Werren and Loehlin, 2009 for line information). Lane 10 is a 100bp ladder (Invitrogen, Thermo Fisher, Waltham, Massachusetts, USA), while Lane 11 is empty.

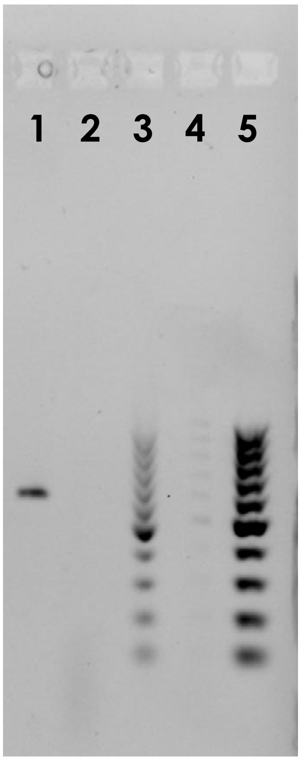

### Figure S.1.1.2 Loss of *Wolbachia* infection, detection via PCR

Detection of *Wolbachia* was carried out according to the PCR protocol of Zhou, Rousset and Neill, 1998. Lane 1 is positive *Wolbachia* control *M. uniraptor* as in Figure S1.1.1. Lane 2 is negative *Wolbachia* control *N. vitripennis* from the AsymCx as in Figure S1.1.1. Lanes 3 and 5 are a 100bp ladder (Invitrogen). Lane 4 is a pooled sample of the general population, showing no products as expected for *Wolbachia*. Similar results were obtained for the S301 inbred line when it was previously used as positive control for other experiments, only to no longer display bands indicative of *Wolbachia* (KBF, unpublished results).

### S1.2 *Wolbachia* contigs from assembly v3.0

*Wolbachia* contigs removed from assembly v3.0, Backbone_1176 and Backbone_1392, available on DANS EASY:

- original/Backbone_1176.fa
- original/Backbone_1392.fa

### S1.3 DAVID Input data

List of gene IDs that were submitted to DAVID for GO term analysis. In the DANS EASY Repository:

- original/Full UniProtKB ID list for DAVID 8247.txt

### S1.4 GenomeScope Results

Figure S1.4.1 GenomeScope results for thelytokous *Trichogramma brassicae* strain S301, based on reads generated in this chapter (BioProject PRJEB35413).

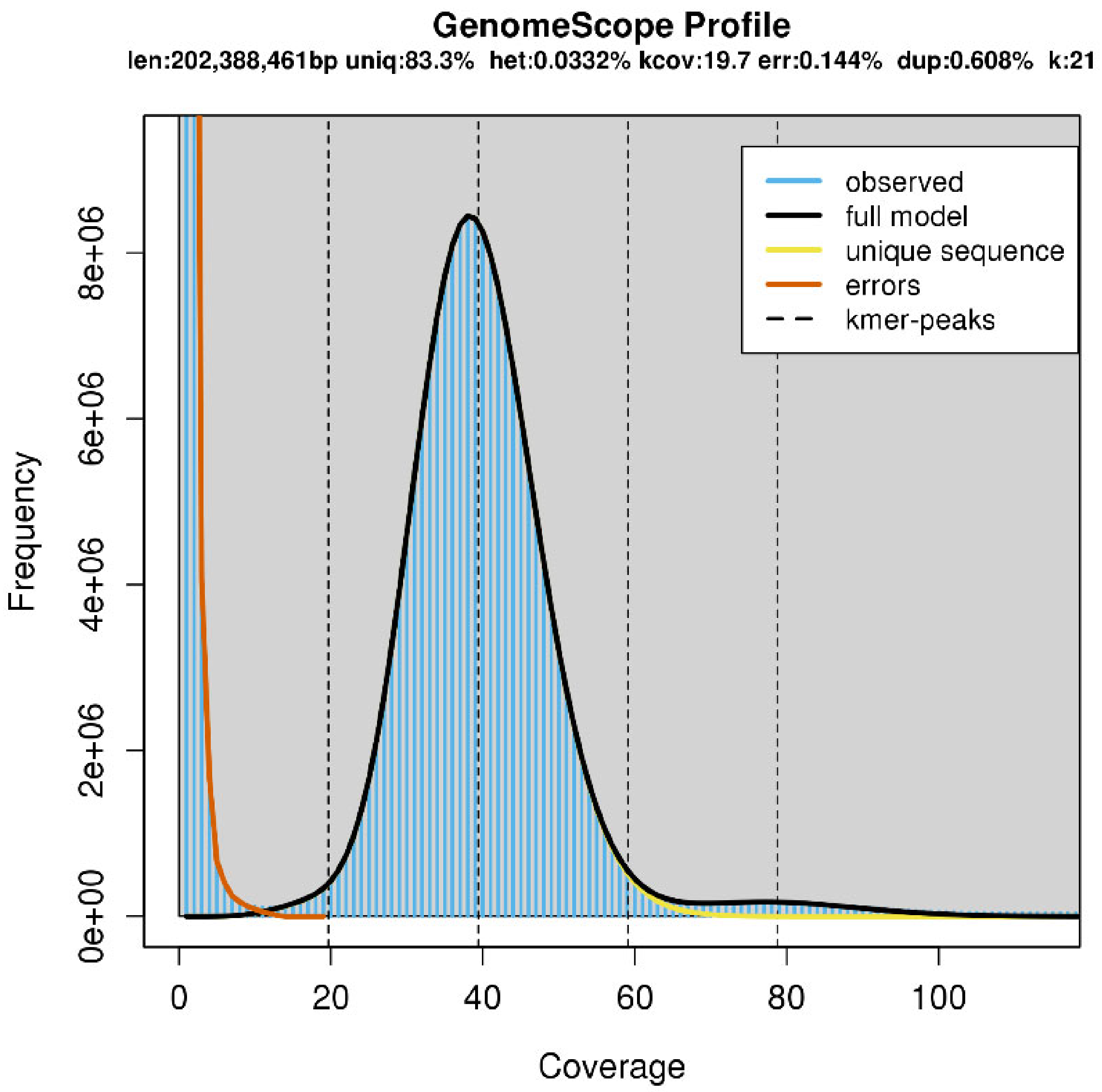

Figure S1.4.2 GenomeScope results for thelytokous strain of *Trichogramma pretiosum*, based on reads from Lindsey et al 2018, NCBI SRA database SRR1191749 (Lindsey et al., 2018).

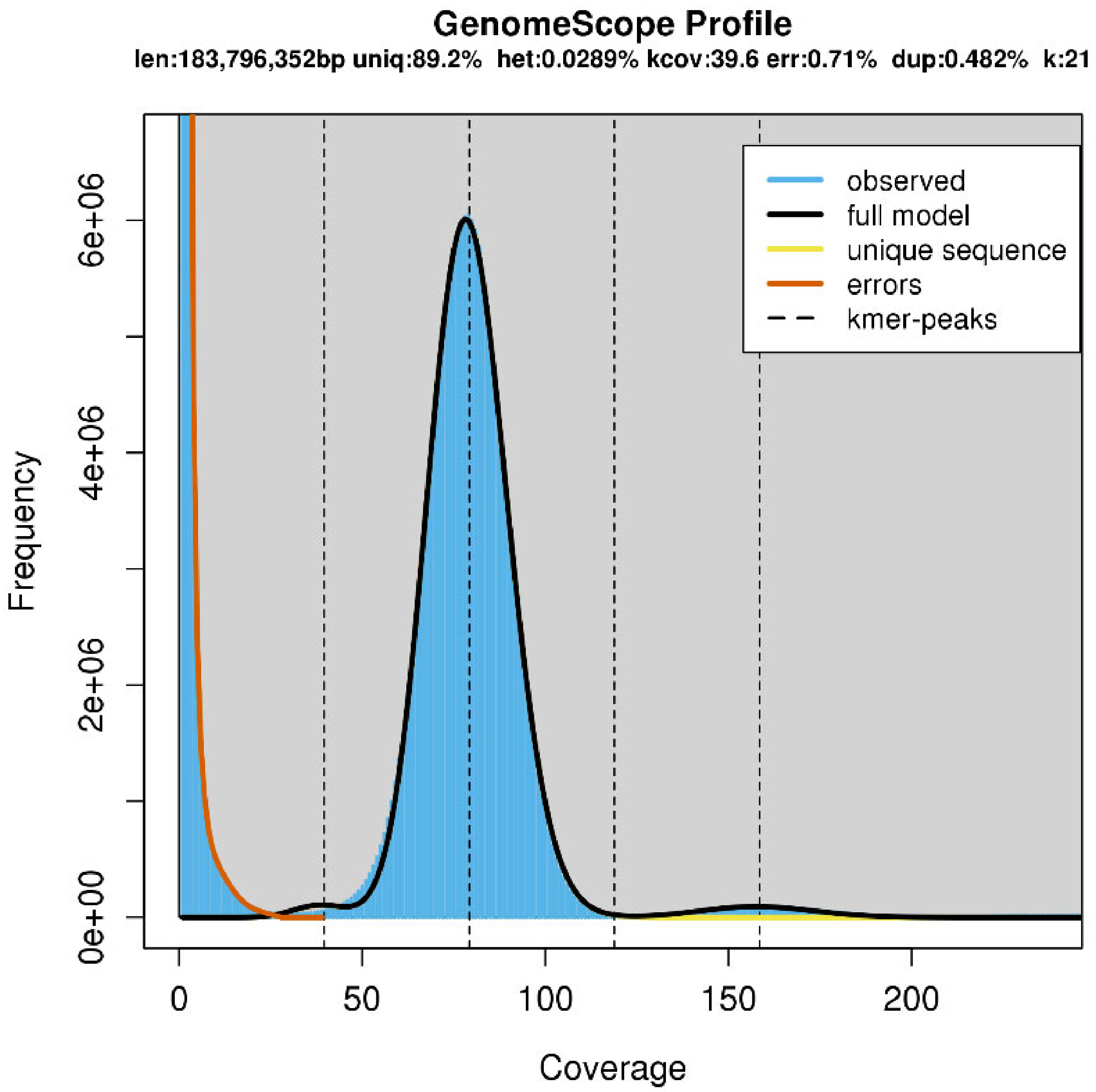

Figure S1.4.3 GenomeScope results for arrhenotokous strain of *Trichogramma pretiosum*, based on reads from Lindsey et al 2018, NCBI SRA database SRR6447489 (Lindsey et al., 2018).

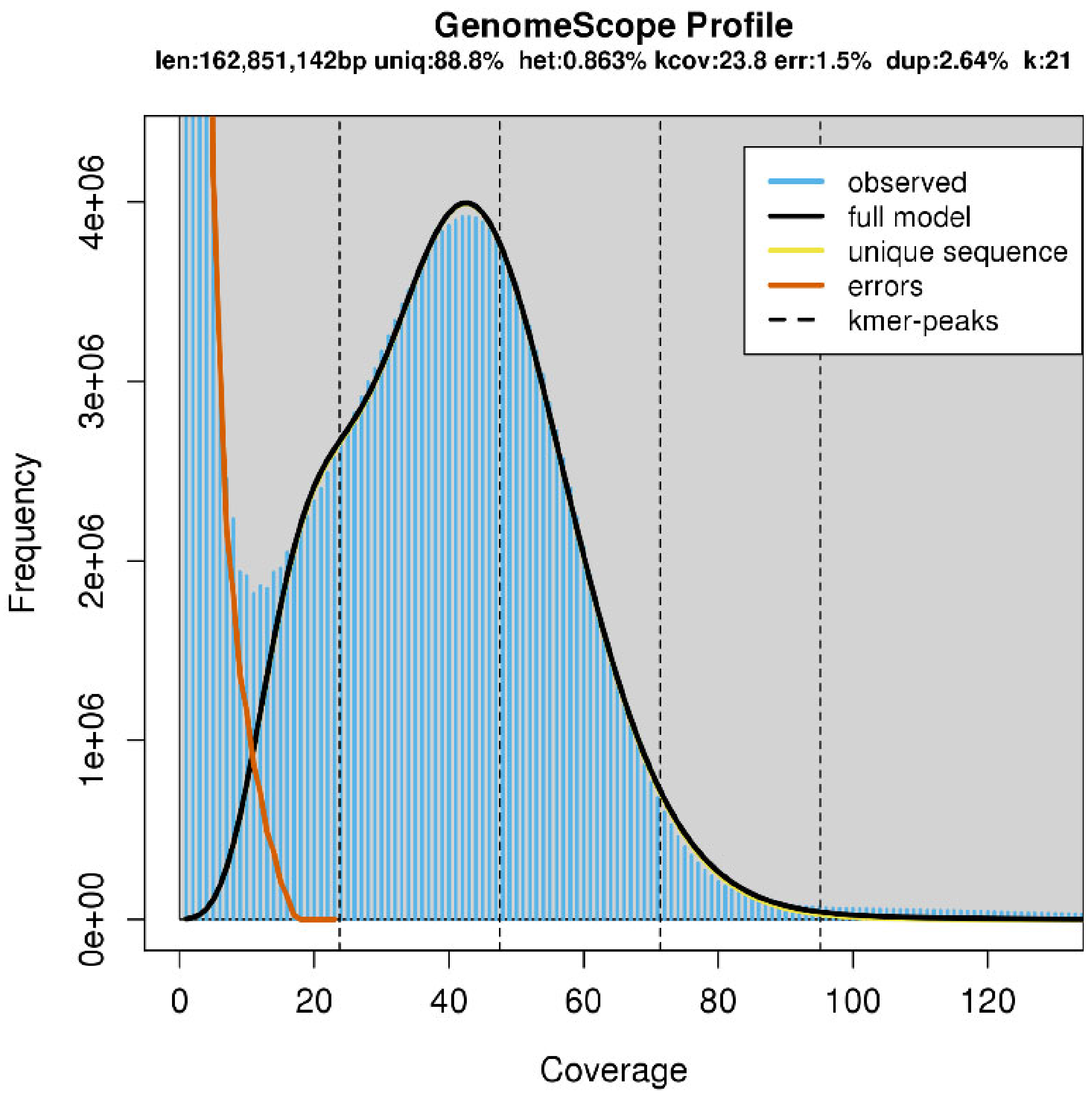

### S1.5 *Trichogramma brassicae* protein set

The official protein set for this annotation, used in ortholog analysis, is available on DANS EASY:

- original/Tricho_Proteins.aa.fasta
